# Fourier transform infrared spectroscopy enables rapid species discrimination across *Malassezia* and strain-level typing in *M. pachydermatis*

**DOI:** 10.1101/2025.09.25.678493

**Authors:** Simon Kurmann, Marco A. Coelho, Marcia David Palma, Leyna Díaz, Gemma Castellá, F. Javier Cabañes, Joseph Heitman, Salomé LeibundGut-Landmann, Francis Muchaamba

## Abstract

1

*Malassezia pachydermatis* is a zoophilic yeast found on the skin and in the outer ear canal of many mammals. It normally maintains a commensal lifestyle but can cause dermatitis and otitis in predisposed hosts, particularly in atopic dogs. *M. pachydermatis* is genetically diverse, with strains clustering into at least three phylogroups based on molecular typing, a pattern we now confirm through whole-genome sequencing (WGS). Accurate species and strain-level identification is essential for understanding its epidemiology, pathogenic potential, and response to treatment. In this study, we established Fourier Transform Infrared (FTIR) spectroscopy as a rapid, cost-effective method for distinguishing *M. pachydermatis* from other *Malassezia* species, including *M. globosa, M. furfur, M. restricta*, and *M. sympodialis*. Within *M. pachydermatis*, FTIR spectroscopy resolved even closely related strains with high accuracy producing clusters congruent with WGS-based phylogeny. The incorporation of an Artificial Neural Network classifier further enhanced the discriminatory power, enabling robust and automated strain assignment. These findings demonstrate the potential of FTIR spectroscopy as a practical tool for large-scale epidemiological surveillance of *M. pachydermatis* and for clinical and veterinary applications where strain-level identification could inform treatment and management of *Malassezia*-associated diseases.

**Importance:** *Malassezia pachydermatis* is a yeast that commonly inhabits the skin and ear canals of mammals but can cause dermatitis and otitis in predisposed hosts, especially dogs with allergies. This species displays substantial genetic diversity, with strains falling into distinct phylogroups that may differ in their biology and clinical significance. Determining these differences has typically required advanced molecular or genomic methods, which can be costly and time-consuming. In this study, we demonstrate that Fourier transform infrared spectroscopy can rapidly and accurately distinguish *M. pachydermatis* from other *Malassezia* species and resolve genetic groups within the species in a way that reflects whole-genome relationships. This capability offers a practical tool for investigating the epidemiology and inter-/intraspecies diversity of *M. pachydermatis* and for guiding targeted management of *Malassezia*-associated diseases in both veterinary and, potentially, human medicine.

## 2 Introduction

*Malassezia* yeasts, a genus of lipophilic basidiomycetes, are found on the skin of warm-blooded animals and humans (1, 2). While usually harmless, they can act as opportunistic pathogens and as such they are implicated in various skin disorders such as dandruff, seborrheic dermatitis, and atopic dermatitis in humans (3), and similar conditions in animals (2). On the other hand, *Malassezia* also undergoes mutual interactions with the host by enhancing the barrier integrity of the skin (4) and by antagonizing skin bacteria to increase colonisation resistance (5, 6).

To date, 18 culturable species of *Malassezia* have been identified (7, 8), with *M. pachydermatis* representing the predominant species in many domestic animals (1, 2, 9, 10) and also in wild animals (11, 12). In healthy dogs, regardless of the breed, *M. pachydermatis* is found in the ear canal and the perioral and interdigital regions (2). Dogs with otitis externa and/or dermatitis have a higher abundance of *M. pachydermatis*. In addition, isolates from diseased dogs were found to exhibit elevated phospholipase A2 activity (13–16), which aligns with the idea that *Malassezia* contributes to pathogenesis by hydrolysing sebum triglycerides into free fatty acids including some that trigger inflammation and skin barrier disruption (17).

Besides its primarily zoophilic nature, *M. pachydermatis* has also zoonotic potential. Rare cases of life-threatening fungemia in humans have been traced back to pet dogs, especially in healthcare settings where hospital workers may act as intermediaries (18). Although *Malassezia* is not typically considered a major public health risk, its presence in hospital environments and the rising antifungal resistance (19) has raised concerns (20).

*M. pachydermatis* displays a high genetic diversity (1, 10, 12, 21, 22). While the clinical significance of this diversity is unclear, it likely affects the fungus’ interaction with the host, reminiscent of what has been described for other opportunistic fungal pathogens such as *Candida albicans* (23). Genetic diversity within *M. pachydermatis* has been explored through multilocus sequence typing (MLST) (1, 10, 24, 25). More recently, *M. pachydermatis* has been sequenced (26). However, only very few complete genomes are currently publicly available (18) for analysing the intraspecies diversity at the genomic level. This scarcity of complete genomic data hinders efforts to fully understand the population structure, evolutionary dynamics, and potential genotype-phenotype associations of *M. pachydermatis*, highlighting the need for additional high-quality genome sequences.

Fourier Transform Infrared (FTIR) spectroscopy offers a compelling alternative for identifying microbial strain variations, with high discriminatory power, as shown for several bacterial species (27, 28) and some fungi, including *Candida* and dermatophytes (29). However, it has not yet been explored for *Malassezia* species. The method is based on quantifying the absorption of infrared light by cellular biomolecules such as nucleic acids, proteins, lipids, and carbohydrates. The interaction of these molecules with infrared light generates a highly specific absorption signature, often referred to as a biochemical fingerprint, characteristic of the organism or strain, and can thus be used for cluster analysis and organism typing (27). Due to its cost- and time-effectiveness, FTIR spectroscopy holds promise for improving the speed and accuracy of strain typing in research and diagnostics.

In this study, we established FTIR spectroscopy as a reliable method for *M. pachydermatis* typing. We show that this method effectively distinguishes *M. pachydermatis* from other *Malassezia* species, discriminates between *M. pachydermatis* strains, and clusters them in a manner comparable to hierarchical clustering based on MLST and whole-genome sequencing (WGS) data. Furthermore, we established a standardized classification system based on the strains’ spectral signatures.

## 3 Materials and Methods

### 3.1 Fungal strains

*M. pachydermatis* (n=30) haploid strains, collected between 1995 and 2018, were selected to represent the species’ genetic diversity (**Table 1**). Strains from other *Malassezia* species (*M. furfur* (n=3), *M. globosa* (n=2), *M. sympodialis* (n=2), and *M. restricta* (n=1)), and from *C. albicans* (n=2), were also included (**Table 2**).

**Table 1.**
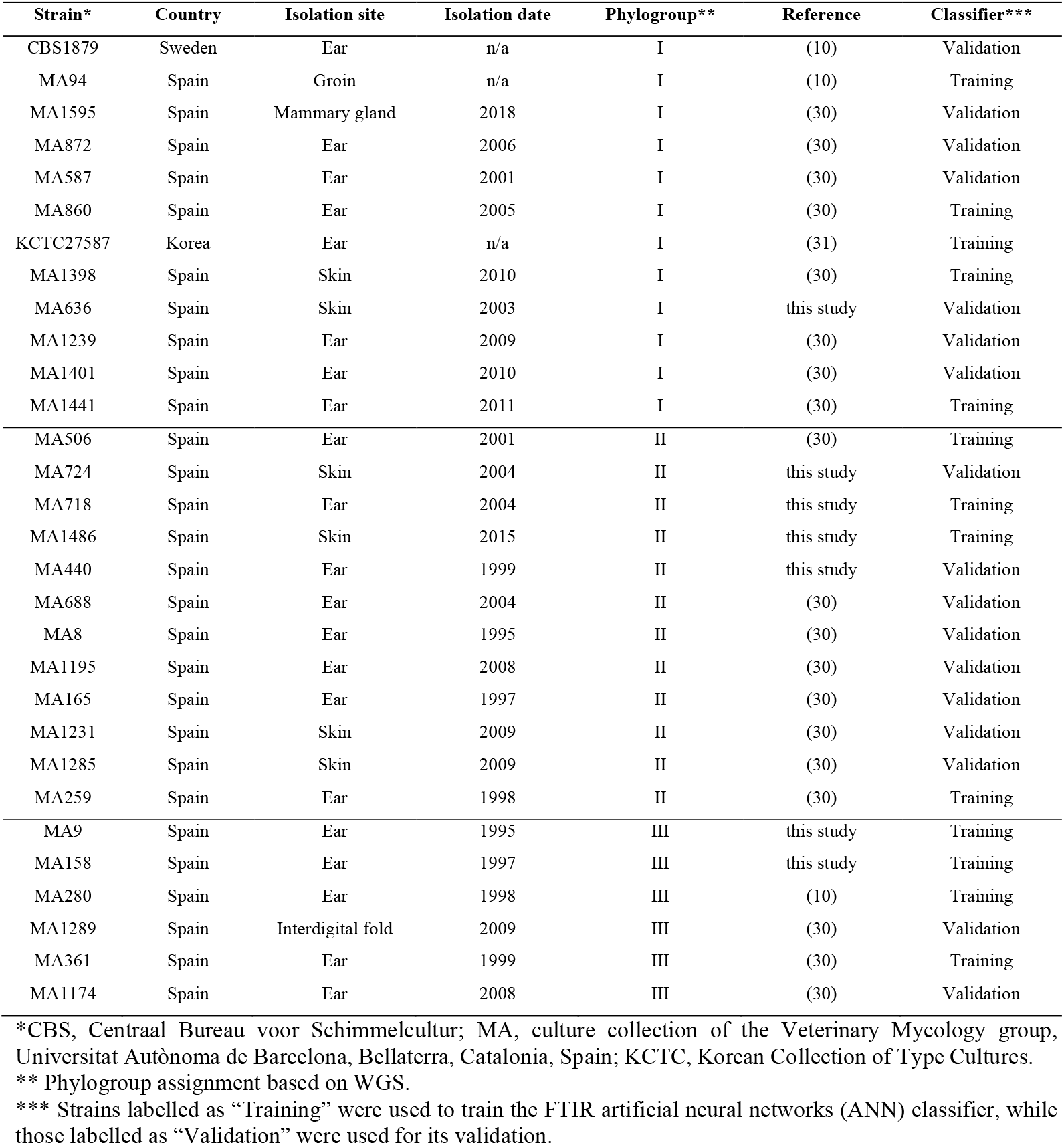
*M. pachydermatis* strains used in this study.

| Strain* | Country | Isolation site | Isolation date | Phylogroup** | Reference | Classifier*** |
| --- | --- | --- | --- | --- | --- | --- |
| CBS1879 | Sweden | Ear | n/a | I | (10) | Validation |
| MA94 | Spain | Groin | n/a | I | (10) | Training |
| MA1595 | Spain | Mammary gland | 2018 | I | (30) | Validation |
| MA872 | Spain | Ear | 2006 | I | (30) | Validation |
| MA587 | Spain | Ear | 2001 | I | (30) | Validation |
| MA860 | Spain | Ear | 2005 | I | (30) | Training |
| KCTC27587 | Korea | Ear | n/a | I | (31) | Training |
| MA1398 | Spain | Skin | 2010 | I | (30) | Training |
| MA636 | Spain | Skin | 2003 | I | this study | Validation |
| MA1239 | Spain | Ear | 2009 | I | (30) | Validation |
| MA1401 | Spain | Ear | 2010 | I | (30) | Validation |
| MA1441 | Spain | Ear | 2011 | I | (30) | Training |
| MA506 | Spain | Ear | 2001 | II | (30) | Training |
| MA724 | Spain | Skin | 2004 | II | this study | Validation |
| MA718 | Spain | Ear | 2004 | II | this study | Training |
| MA1486 | Spain | Skin | 2015 | II | this study | Training |
| MA440 | Spain | Ear | 1999 | II | this study | Validation |
| MA688 | Spain | Ear | 2004 | II | (30) | Validation |
| MA8 | Spain | Ear | 1995 | II | (30) | Validation |
| MA1195 | Spain | Ear | 2008 | II | (30) | Validation |
| MA165 | Spain | Ear | 1997 | II | (30) | Validation |
| MA1231 | Spain | Skin | 2009 | II | (30) | Validation |
| MA1285 | Spain | Skin | 2009 | II | (30) | Validation |
| MA259 | Spain | Ear | 1998 | II | (30) | Training |
| MA9 | Spain | Ear | 1995 | III | this study | Training |
| MA158 | Spain | Ear | 1997 | III | this study | Training |
| MA280 | Spain | Ear | 1998 | III | (10) | Training |
| MA1289 | Spain | Interdigital fold | 2009 | III | (30) | Validation |
| MA361 | Spain | Ear | 1999 | III | (30) | Training |
| MA1174 | Spain | Ear | 2008 | III | (30) | Validation |

\*CBS, Centraal Bureau voor Schimmelcultuur; MA, culture collection of the Veterinary Mycology group, Universitat Autònoma de Barcelona, Bellaterra, Catalonia, Spain; KCTC, Korean Collection of Type Cultures.
\*\* Phylogroup assignment based on WGS.
\*\*\* Strains labelled as “Training” were used to train the FTIR artificial neural networks (ANN) classifier, while those labelled as “Validation” were used for its validation.

**Table 2.** Other yeast strains used in this study.

| Species | Strain ID | Host species | Reference |
| --- | --- | --- | --- |
| *M. furfur* | CBS1878 | human | (25) |
| *M. furfur* | CD866 | human | (32) |
| *M. furfur* | CBS9595 | human | (32) |
| *M. globosa* | CBS7966 | human | (25) |
| *M. globosa* | GB_AD72_270223 | human | this study |
| *M. sympodialis* | ATCC 42132 | human | (33) |
| *M. sympodialis* | 10106 (KS016) | human | (34) |
| *M. restricta* | RS HS_19 | human | this study |
| *C. albicans* | SC5314 | human | (35) |
| *C. albicans* | 101 | human | (23) |

### 3.2 Culture conditions

All *Malassezia* strains were preserved in glycerol at -80°C and revived by transferring an aliquot on modified Dixon (mDixon) agar (36 g malt extract (Sigma), 20 g desiccated oxbile (Sigma), 6 g bacto peptone (Becton Dickinson), 10 mL Tween-40 (Sigma), 2 mL pure glycerol (Sigma), 2 mL oleic acid (Sigma, stored at -20°C), and 15 g agar (Sigma) per Liter, pH adjusted to 6.0 using HCl) and incubating the plates for 2-5 days at 30°C. *C. albicans* strains were grown on YPD agar (20 g D-glucose (Cayman Chemicals), 10 g bacto yeast extract (Becton Dickinson), 20 g bacto peptone, 20 g agar per Liter) for 24 hours at 30°C. For FTIR analysis, individual fungal colonies were sub-cultured on fresh mDixon agar for exactly 48 hours at 30°C. *M. restricta*, was incubated for 7 days due to its slow growth rate.

### 3.3 FTIR Spectroscopy

FTIR spectroscopy analysis was performed using the IR Biotyper^®^ (Bruker Daltonics), following the manufacturer’s instructions with minor modifications. For sample preparation, two 1 µL inoculation loops of biomass were transferred into 70 µL of deionized water in a 1.5 mL tube containing metal rods (Bruker Daltonics). The suspension was homogenized by shaking on a MSC 100 Thermo Shaker (Labgene Scientific) at 1400 rpm and room temperature for 10 minutes. Subsequently, 70 µL of 70% (v/v) ethanol was added, followed by an additional 5 minutes of shaking. For initial colony resuspension, the reverse was also tried, with ethanol added first, followed by water. In this case, two 1 µL inoculation loops of biomass were transferred into 70 µL of 70% (v/v) ethanol in a 1.5 mL tube containing metal rods, homogenized by shaking at 1400 rpm for 10 minutes. Then, 70 µL of deionized water was added, and the suspension was shaken for an additional 5 minutes. Next, 15 µL of the resulting suspension was spotted in triplicates on a silicon-based 96-sample plate (Bruker Daltonics) and let dry at 37°C. 10 μL of the reconstituted Infrared Test Standards (IRTS 1 and IRTS 2) were also spotted in duplicate onto the IR Biotyper^®^ target and let dry as described for the samples. Spectra acquisition was performed using the IR Biotyper^®^ system (OPUS software v8.2; Bruker Optics) with default analysis settings. Three independent biological repeats (independent colonies from independent plates), with at least three technical replicates each, were conducted for each strain.

Data analysis was performed using IR Biotyper^®^ Client Software v4.0 (Bruker Daltonics). Only spectra that passed the software-intrinsic quality control were further analysed. Spectral windows corresponding to proteins, lipids, polysaccharides, or their combinations were evaluated to identify the most discriminatory region. Subsequently, spectral splicing methods covering the range characteristic for polysaccharides (1300–800 cm^−1^) were applied, unless specified differently. The spectra were pre-processed by calculating the second derivative, splicing them according to regions, and applying vector normalization. Clustering of spectra was performed using Hierarchical Cluster Analysis (HCA), Principal Component Analysis (PCA), and Linear Discriminant Analysis (LDA) for both supervised and unsupervised classification. Euclidean distance and average linkage clustering methods were applied to define clusters, using the determined Cut-Off Values (see below). Results were visualized through dendrograms, distance matrices, and 2-D and 3-D scatter plots.

Starting from the Cut-Off values automatically generated by the software, an optimization procedure was conducted to determine the optimal values for *M. pachydermatis*, tailored to our laboratory-specific conditions. This optimization was informed by the genetic relatedness of the strains, as determined by MLST and WGS (see below).

### 3.4 Genomic DNA extraction, genome sequencing and assembly

Genomic DNA was extracted from selected *M. pachydermatis* strains using a phenol:chloroform-based protocol as previously described (36). DNA concentrations were quantified with the Qubit dsDNA Assay Kit (Invitrogen) using a Qubit fluorometer. Whole-genome sequencing was performed at the Duke University Sequencing and Genomic Technologies (DUSGT) Core Facility. Libraries were prepared with the Kapa HyperPlus kit and sequenced on the Illumina NovaSeq platform to generate 2 × 150 bp paired-end reads. Draft genome assemblies were produced with SPAdes v3.15.3 (37), using default parameters and retaining contigs longer than 500 bp.

### 3.5 Phylogenetic and ANI analyses

Phylogenetic relationships were inferred from both genome-wide SNP data and MLST markers. For the whole-genome analysis, high-confidence single nucleotide variants (SNVs) were called with Snippy v4.6.0, using the *M. pachydermatis* KCTC27587 genome as reference and the following parameters: *--cpus 10 --unmapped --mincov 10 --minfrac 0*.*9*. The resulting core genome alignment (30 sequences, 373,384 sites) served as input for maximum likelihood (ML) tree reconstruction with IQ-TREE v2.3.6, applying the following parameters:

*-m MFP -msub nuclear --bnni -B 1000 -alrt 1000*, which enables model selection, nuclear-specific substitution models, and branch support via 1,000 ultrafast bootstrap (UFboot) replicates and SH-aLRT tests.

MLST-based phylogenies were reconstructed from both concatenated and single-locus alignments. Sequences of four loci previously shown to differentiate *M. pachydermatis* strains—large subunit (LSU) and internal transcribed spacer (ITS) rRNA regions, *CHS2* (chitin synthase 2), and *TUB2* (β-tubulin)—were retrieved from each genome assembly via BLASTN searches using one deposited reference sequence per marker as query (10). For each locus, the top match per strain was extracted with *samtools faidx*, and reverse-strand hits were reoriented. The retrieved sequences were aligned with MAFFT v7.310 (*--adjustdirection*, FFT-NS-2 mode) and trimmed to the defined amplicon boundaries established in the original MLST scheme (10). In the concatenated analysis, the four trimmed alignments (30 sequences, 4 partitions, 5,434 sites) were merged into a partitioned supermatrix and analysed with IQ-TREE v2.3.6 using the same ML parameters described above, with the *-p* option to define partitions. In parallel, single-gene ML trees were constructed to assess the phylogenetic resolution of each marker and to assign allele types. Each gene alignment was supplemented with published reference sequences representing known MLST alleles (10, 12, 30) (**Supplementary Table S1**), and individual trees were inferred with IQ-TREE under the best-fit substitution model selected per gene, with 10,000 UFboot and SH-aLRT replicates to improve branch support resolution for the shorter alignments.

Pairwise average nucleotide identity (ANI) values were computed with OrthoANIu (38) (**Supplementary Table S2-S3**). Custom Python scripts transformed ANI values into distance scores (100 − ANI), applied average linkage clustering, and generated a Newick-format dendrogram. Group-wise ANI statistics (minimum, maximum, mean, and number of comparisons) were derived from a strain-to-clade mapping file. All phylogenetic trees and dendrograms were visualized with iTOL v7.

### 3.6 Comparison of FTIR and WGS-based clusters

Three different metrics were employed to compare clustering concordance between FTIR- and WGS-based approaches, calculated using an online tool (http://www.comparingpartitions.info/) with 95% confidence intervals (CIs) (39). The Simpson’s Index of Diversity (SID) was calculated separately for FTIR- and WGS-based clusterings to evaluate the diversity of clusters identified by each method. The Adjusted Rand Index (ARI) measured the degree of agreement between clustering methods while correcting for chance, with values closer to 1 indicating stronger concordance (39–41). Finally, the Adjusted Wallace Coefficient (AWC) quantified directional congruence between clustering methods. Specifically, the AWC quantifies the probability that two strains clustered together by one method (e.g., WGS) are also clustered together by the other (e.g., FTIR spectroscopy), and vice versa, with a value of 1 indicating complete agreement between two methods. Optimal phylogroup-specific FTIR cut-off values were defined by maximizing the ARI between FTIR- and ANI-based clustering.

### 3.7 Development of an automated classifier for *M. pachydermatis*

To enable the automated classification of *M. pachydermatis* strains based on FTIR spectra, we developed a classifier using artificial neural networks (ANN) integrated within the IR Biotyper^®^ software. This classifier was designed to rapidly identify unknown samples based on spectral data, using marker models derived from a set of training spectra to categorize strains into three distinct *M. pachydermatis* phylogroups (I, II, and III). For classifier development, strains were divided into training and validation sets (**Table 1**). The training set, comprising five phylogroup I strains and four strains each from phylogroups II and III, was used to develop the classifier (n=9 spectra per strain). The remaining 17 strains (n=9 spectra per strain) formed the validation set for evaluating classifier performance. To ensure comprehensive representation of spectral variability, including outliers, training strains were selected based on deviation plots of the individual vector-normalized average spectrum of each strain within its respective phylogroup (**Supplementary Figure S1A-C**). Data from three biological replicates were used per strain to enhance the robustness and reproducibility of the classifier. Principal components (PCs) for training were selected using an overall vector-normalized spectral deviation plot of all three phylogroups (**Supplementary Figure S1D**), allowing exclusion of components that primarily contributed noise or non-informative variance. Training cycle parameters were optimized to avoid overfitting, aiming to produce a classifier applicable to real-world data. Classifier performance was evaluated using stratified 4-fold cross-validation, with a confusion matrix generated to assess classification accuracy.

The classification results were reported using a scoring system with colour coding to indicate confidence levels. A “green” score denoted a high-confidence match, where the sample spectrum fell within the spectral space of the training set (outlier value <1.0). A “yellow” score indicated moderate confidence, corresponding to spectra on the periphery of the spectral space of the training set (outlier value between 1.0 and 2.0). “Red” was assigned when the sample spectrum was outside the training set spectral space (outlier value >2.0), categorizing it as “no classification possible.” Misclassifications were defined as incorrect classifications assigned green or yellow scores. Classifier performance was evaluated based on accuracy (the proportion of correctly classified spectra), error rate (misclassified spectra/total spectra), sensitivity (the proportion of target strains correctly identified), specificity (the proportion of non-target strains correctly excluded), and positive and negative predictive values (**Supplementary Table S4)**. Additionally, classifier versatility was assessed using spectra from non-*M. pachydermatis* strains to evaluate its capacity to exclude unrelated species and its potential for broader application.

### 3.8 Data availability statement

All other raw data linked to this study will be made publicly available at zenodo.org upon acceptance of the manuscript (doi will be provided). Raw sequencing reads, and genome assemblies were deposited in the DDBJ/EMBL/GenBank databases under BioProject PRJNA1330337 (detailed in **Supplementary Table S5**).

## 4 Results

### 4.1 Phylogeny of *M. pachydermatis* strains based on MLST and WGS

To investigate the genomic diversity of the 30 *M. pachydermatis* strains included in this study, we first performed MLST analysis with four loci—*CHS2*, LSU (D1/D2 domain of the large subunit rRNA), ITS, and *TUB2*—previously shown to differentiate *M. pachydermatis* (10), using sequences extracted from newly generated draft genome assemblies. The concatenated MLST phylogeny resolved the strains into three well-supported phylogroups, here designated I, II, and III (**Figure 1A; Supplementary Figure S2**). Among the single-gene trees, the LSU provided the clearest separation of these groups, while ITS and *TUB2* showed higher levels of intra-group polymorphism. In addition to confirming previously defined allele types, our analysis identified novel genotypes at the ITS (type XIX in strains MA1486, MA1195, MA724, MA506; and type XX in strains MA158 and MA1174) (**Supplementary Figure S2**), thereby expanding the documented allelic diversity of this species.

**Figure 1.**
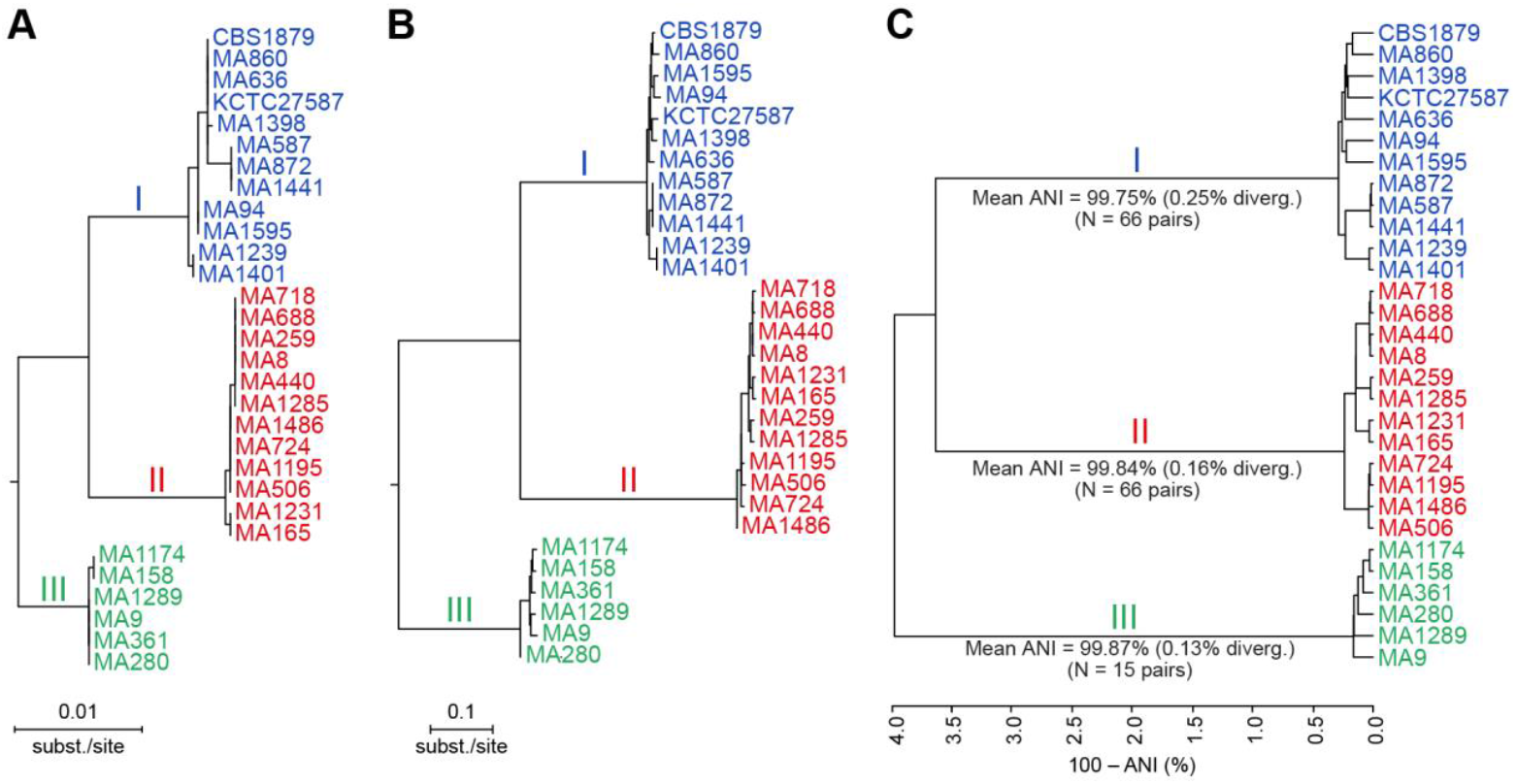
Phylogenetic relationships and sequence divergence among *M. pachydermatis* strains. **(A)** Maximum likelihood (ML) phylogeny based on a concatenated multilocus sequence typing (MLST) alignment comprising 5,434 nucleotide sites across four loci: ITS1, LSU, *CHS2*, and *TUB2*. IQ-TREE was run with partitioned models. Branch support (100% ultrafast bootstrap and SH-aLRT) was confirmed only for the three major phylogroups; internal nodes received low support and are not annotated for simplicity. **(B)** ML phylogeny inferred from 373,384 genome-wide single nucleotide variants, based on the core genome alignment. Tree reconstruction was performed under the best-fit substitution model TVM+F+ASC+R2, selected by BIC. All branches received 100% support from both ultrafast bootstrap and SH-aLRT tests. Both phylogenetic trees (A and B) were midpoint rooted. **(C)** UPGMA dendrogram based on pairwise sequence divergence, calculated as 100 − Average Nucleotide Identity (ANI), with values computed by OrthoANIu. Mean ANI, corresponding average divergence, and the number of pairwise comparisons are indicated for each phylogroup. Strains are coloured by phylogroup: group I (blue, n= 12), group II (red, n = 12), and group III (green, n = 6).

We next reconstructed phylogenetic relationships using WGS data to achieve higher resolution. A maximum-likelihood tree based on 373,384 high-confidence SNVs from the core genome confirmed the assignment of all strains to the three phylogroups initially delineated by MLST, with strong branch support (**Figure 1B**). All major nodes received 100% support in both ultrafast bootstrap and SH-aLRT tests, showing the robustness of the WGS-based classification.

Pairwise ANI values further highlighted differences in intra- and inter-clade diversity (**Figure 1C; Supplementary Table S2**). Phylogroup I (n=12) exhibited the greatest heterogeneity, with a mean ANI of 99.75% corresponding to 0.25% average genomic divergence. Phylogroup II (n=12) showed intermediate variability (mean ANI 99.84%; divergence 0.16%), whereas phylogroup III (n = 6) was the most homogeneous group (mean ANI 99.87%; divergence 0.13%). Comparative divergence between groups revealed that phylogroup I was genetically closer to phylogroup II (3.63%) than to phylogroup III (3.68%), while phylogroups II and III were the most divergent pair (4.26%) (**Supplementary Table S3**).

Together, these results demonstrate that *M. pachydermatis* is structured into three well-supported genomic lineages, identifiable by both MLST and WGS. While MLST provides a useful framework for initial strain classification, WGS offers greater resolution, enabling robust phylogenetic inference and quantitative assessments of genomic divergence. These findings establish a phylogenetic framework for *M. pachydermatis*, providing the basis for future studies to explore how lineage-specific genomic variation may influence host interactions, pathogenic potential, or ecological adaptation.

### 4.2 Establishing of a working protocol for analysing *Malassezia* by FTIR

While WGS provides high-resolution insights, its routine use remains limited, therefore, we explored FTIR spectroscopy as a high-throughput and cost-effective alternative for strain typing. For establishing a standardized protocol for FTIR analysis of *Malassezia* spp. using the IR Biotyper^®^, we first developed a reliable sample preparation protocol that ensured that all strains consistently passed the initial quality control checks of the IR Biotyper^®^ software. To overcome the limited ability to resuspend *Malassezia* colony material in ethanol (**Supplementary Figure S3**), the fungal colony material was first resuspended in water, followed by addition of an equal volume of 70% ethanol. This procedure contrasted with the manufacturer’s ethanol-first method, which, even after extended vortexing (>15 minutes), failed to yield homogeneous suspensions of colony material and resulted in poor-quality spectra that frequently failed the quality control check. Our modified “water-first, then ethanol” approach yielded FTIR spectra of high signal quality that consistently passed the IR Biotyper^®^ software’s quality control checks. The resulting spectra were highly reproducible across both technical and biological replicates of the strains. This optimized protocol was thus adopted as the default method for the entire study.

To evaluate reproducibility of the method, FTIR spectra of three *M. pachydermatis* strains were acquired on three different days with three technical replicates each. Both the technical and biological replicates of these strains clustered tightly for each strain (**Figure 2**), highlighting the reproducibility and reliability of the protocol.

**Figure 2.**
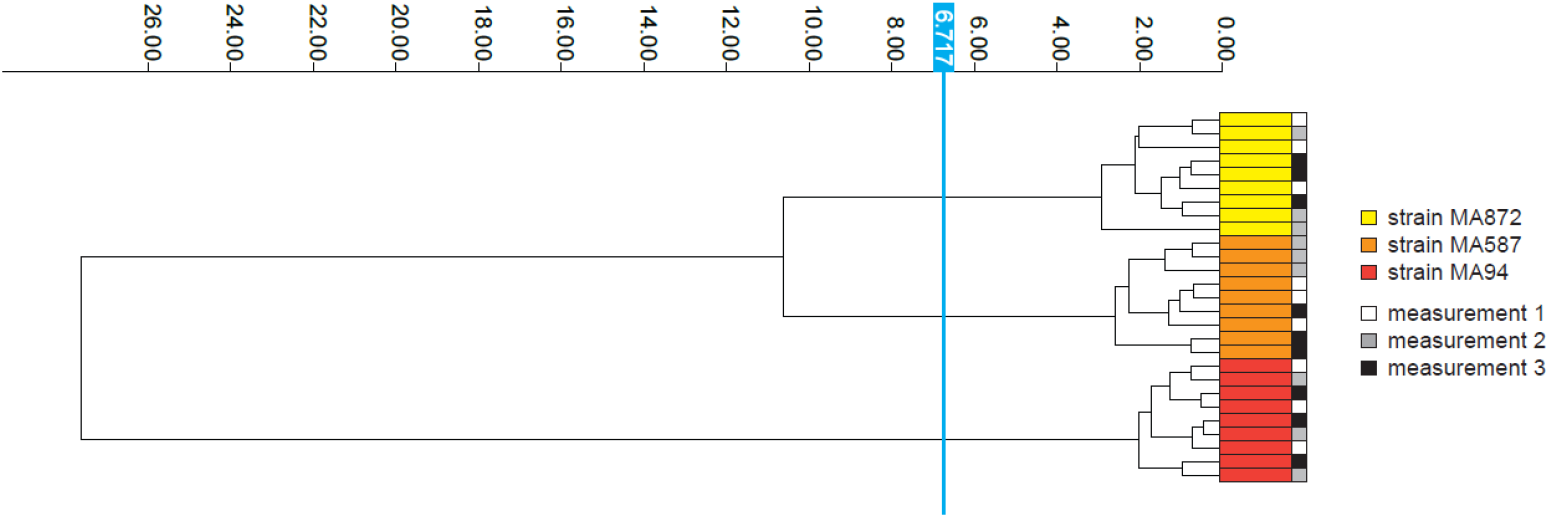
FTIR is highly reproducible across technical and biological replicates. Dendrogram generated from clustering FTIR spectra of *M. pachydermatis* strains (MA872, MA587, MA94) to assess reproducibility across technical and biological replicates. Spectra are color-coded by strain and acquisition date, with each row representing a technical replicate derived from three independent experiments conducted on three separate days. The vertical blue line denotes the automatically calculated cut-off value (6.717) for clustering. Dimensionality reduction: LDA (2 LDs of 10 principal components (PCs); 95.7% variance).

Comparison of the spectral ranges corresponding to polysaccharides (1300-800 cm-^1^), proteins (1800-1500 cm-^1^), lipids (3000-2800 cm-^1^ and 1500-1400 cm-^1^), and polysaccharide-protein combinations (1800 -900 cm-^1^), revealed that the splicing method for the polysaccharide region exhibited the highest discriminatory power (**Supplementary Figure S4**). Therefore, all subsequent analyses focused on the polysaccharide spectral region.

### 4.3 FTIR spectroscopy discriminates between different *Malassezia* spp

After establishing a robust protocol for applying FTIR spectroscopy to *Malassezia*, we investigated the discriminatory power of the method at both the genus and species level. For this, we compared the FITR spectra of several different *Malassezia* spp. We also included two

*C. albicans* strains in the analysis, which clustered distantly from *Malassezia* spp. (**Figure 3**). Within the *Malassezia* genus, individual species clustered distinctly from each other (**Figure 3**), confirming the potential of FTIR for species-level differentiation. Within species, different strains were clearly segregated from each other, with biological and technical replicates always clustering most closely (**Figure 3**).

**Figure 3.**
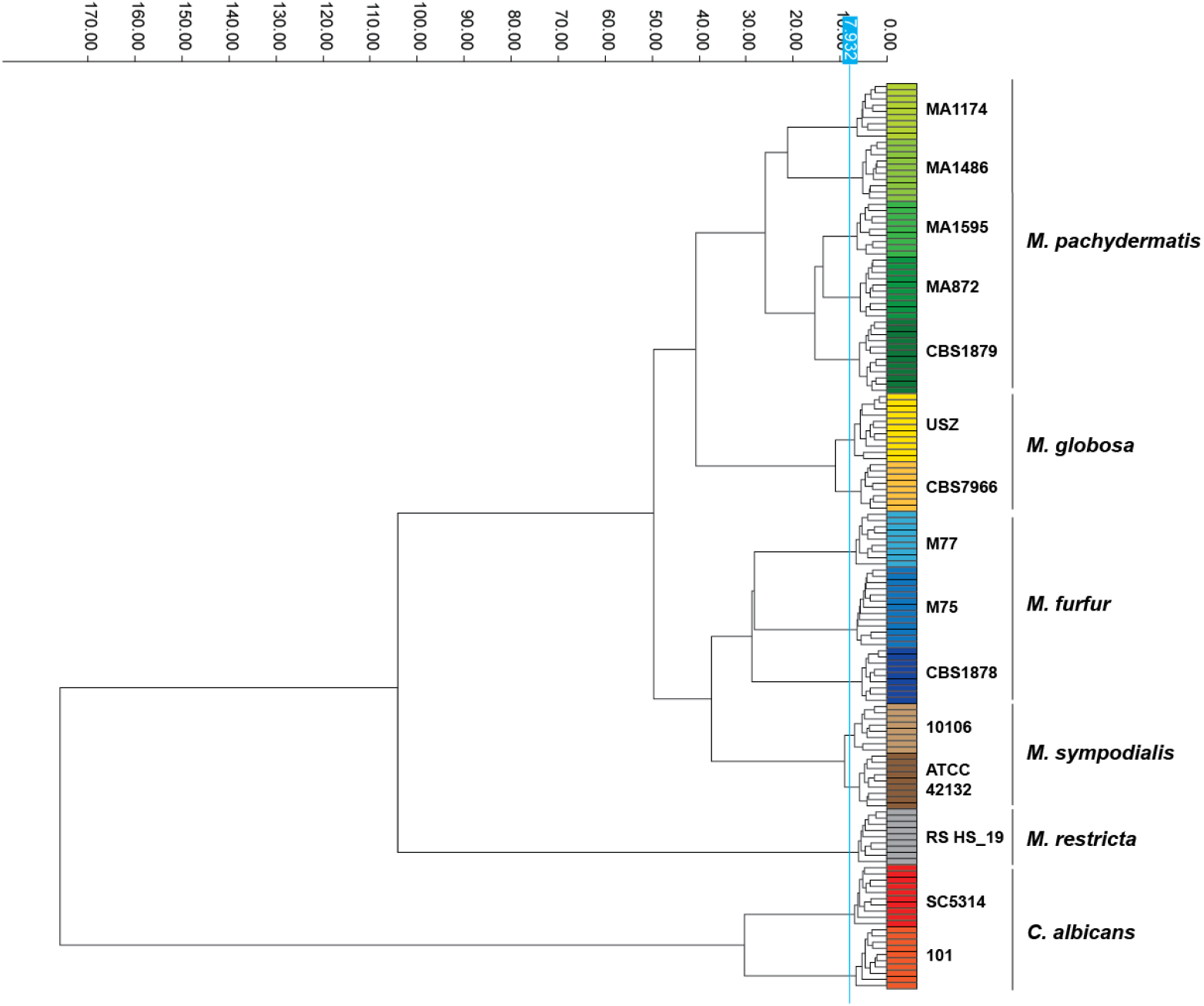
FTIR spectroscopy discriminates between different *Malassezia* spp. Dendrogram generated from clustering FTIR spectra of *Malassezia* spp. and *Candida albicans* to assess species differentiation. Spectra are color-coded by strain and species, with each row representing a technical replicate pooled from three independent biological repeats. The vertical blue line denotes the automatically calculated cut-off value (7.932) for clustering. Dimensionality reduction: LDA (14 LDs of 30 principal components (PCs); 99.8% variance).

### 4.4 FTIR spectroscopy achieves strain-level resolution in *Malassezia pachydermatis*

Next, we evaluated the capacity of FTIR spectroscopy to discriminate among *M. pachydermatis* strains, where a larger number of isolates was available, and to assess its effectiveness in differentiating strains belonging to different phylogroups. Analysing the FTIR spectra of the 30 strains (**Table 1**) using linear discriminant analysis (LDA) successfully grouped them into three clusters corresponding to phylogroups I, II and III (**Figure 4A, Supplementary Figure S5**). Phylogroups I and II were well separated (**Figure 4B, Supplementary Figure S6**), while phylogroup III exhibited greater internal variability (**Figure 4, Figure 5**). Within phylogroup III, two strains formed a cluster clearly separate from clusters I and II, while the remaining four strains (MA9, MA361, MA1289, and MA1174) were found in closer proximity to phylogroup I or II strains (**Figure 4A, Supplementary Figure S5**).

**Figure 4.**
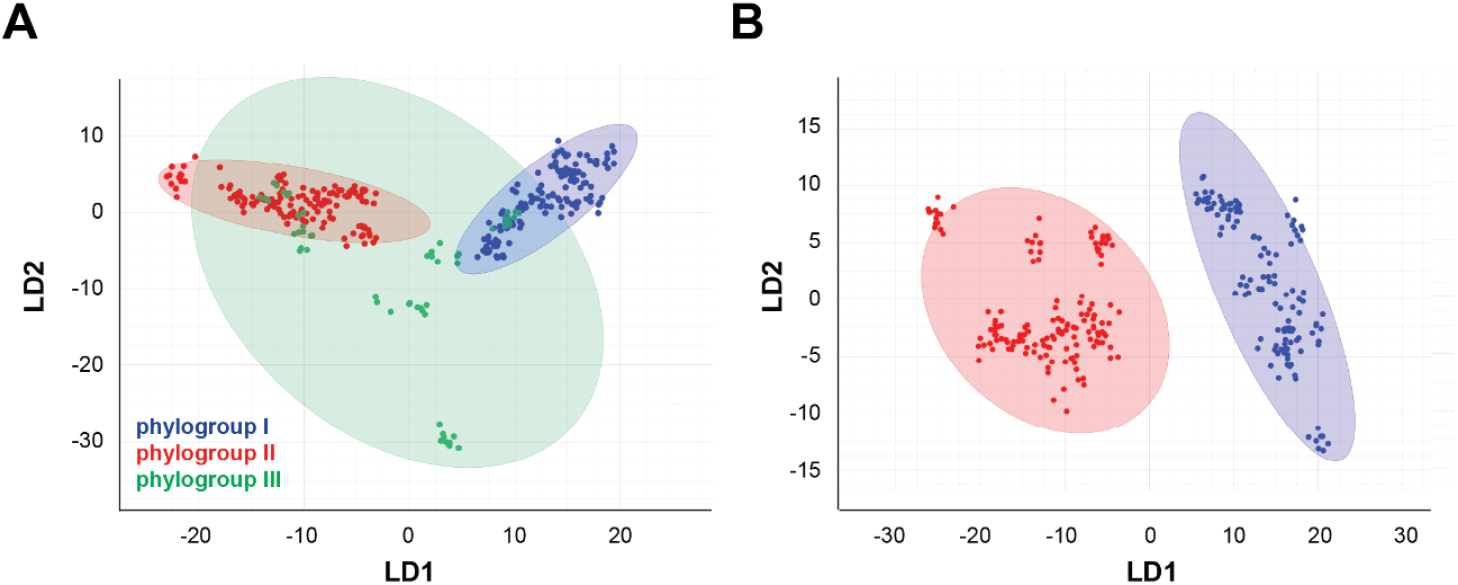
FTIR spectroscopy achieves strain-level resolution in *M. pachydermatis*. Scatter plots showing the distribution of *M. pachydermatis* phylogroups I (n=12), II (n=12), and III (n=6) strains (**A**), and phylogroups I (n=12) and II (n=12) strains (**B**) within the FTIR spectral space. Each strain is represented by nine spectra (•) pooled from three independent biological replicates with three technical replicates each. Data were processed using Linear Discriminant Analysis (LDA) based on 30 principal components. The diagram displays the first two LD axes, with spectra color-coded by phylogroup (I: blue, II: red, III: green).

**Figure 5.**
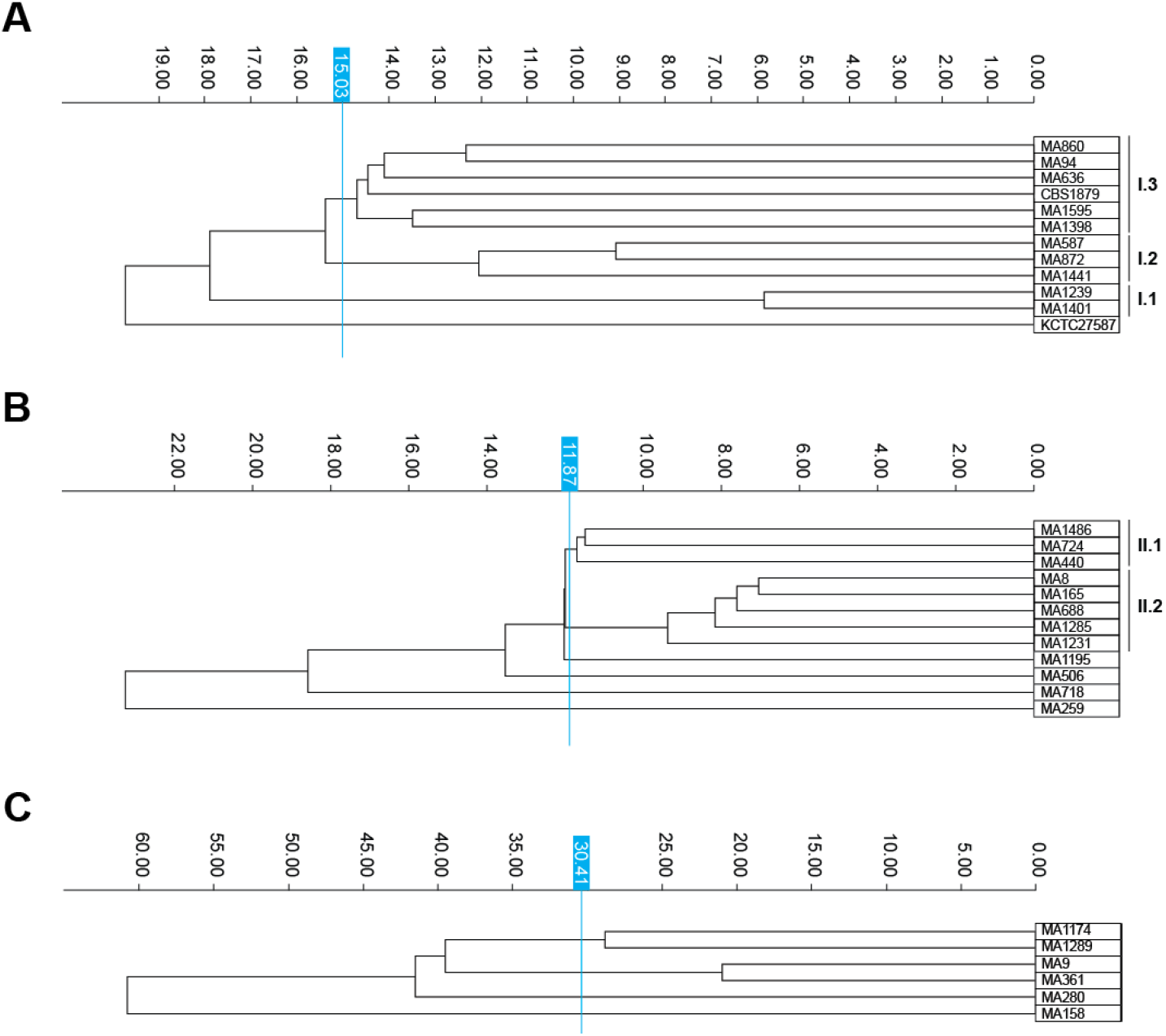
Subclusters in phylogroups I, II, and III based on FTIR spectral profiles. Dendrogram generated from clustering FTIR spectra of *M. pachydermatis* strains of phylogroups I, II, and III separately to assess intra-phylogroup variability. Each row represents the average spectrum of a strain, calculated from at least nine individual absorption spectra (three independent biological replicates with three technical replicates each per strain). The vertical blue lines denote the cut-off value for clustering. Dimensionality reduction: (**A**) phylogroup I: LDA (11 LDs of 30 principal components (PCs); 98.7% variance), (**B**) phylogroup II: LDA (11 LDs of 30 principal components (PCs); 99.2% variance), and (**C**) phylogroup III: LDA (5 LDs of 30 principal components (PCs); 99.7% variance). Subclusters are indicated to the right.

A more detailed analysis of the FTIR spectra, conducted separately for each phylogroup, identified distinct clustering patterns (**Figure 5, Supplementary Figure S7**). In phylogroup I, three subclusters were identified: subcluster I.1 included strains MA1239 and MA1401; subcluster I.2 comprised strains MA587, MA872, and MA1441; and subcluster I.3 contained strains MA94, MA636, MA860, MA1398, MA1595, and CBS1879. Strain KCTC27587 was positioned more distantly from the other strains of this phylogroup. Pairwise relatedness analysis further showed that strains MA1239 and MA1401 were the most closely related pair (spectral distance 5.85), followed by MA587 and MA872 (spectral distance 9.08). In phylogroup II, two subclusters were observed: subcluster II.1 included strains MA440, MA724, and MA1486, while subcluster II.2 comprised strains MA8, MA165, MA688, MA1231, and MA1285. Four strains (MA259, MA506, MA718, and MA1195) did not cluster within either of these subgroups. Phylogroup III exhibited the highest intra-group variability, with no clear subclusters observed (**Supplementary Figure S7C**).

### 4.5 Comparison of the strain clustering based on FTIR spectroscopy vs. WGS-based clustering

We next compared FTIR clustering to WGS, the gold standard for microbial classification, using three metrics: Simpson’s index of diversity (SID), adjusted Rand index (ARI), and adjusted Wallace coefficient (AWC). Overall, WGS exhibited slightly higher diversity (SID: 0.662, 95% CI: 0.604–0.720) compared to FTIR (SID: 0.605, 95% CI: 0.527–0.682). FTIR reproduced the WGS-based clustering of phylogroup I (n=12) and II (n=12) strains with perfect agreement, as reflected by ARI and AWC values of 1.00. Notably, FTIR clearly distinguished phylogroups I and II with high confidence, despite these groups being more similar to each other based on ANI (mean divergence: I vs. II = 3.63%, I vs. III = 3.68%, II vs. III = 4.26%). Furthermore, phylogroup III, which showed the least diversity by WGS, exhibited the highest diversity in FTIR profiles, suggesting that FTIR captures phenotypic variation not evident in genomic comparisons. In the case of phylogroup III strains, FTIR misclassified three strains, resulting in an overall ARI of 0.768 when considering all studied strains (n=30). Despite this misclassification, the AWC indicated that FTIR clusters were highly predictive of WGS clusters (AWC: 0.876, 95% CI: 0.824–0.929), highlighting FTIR’s potential as a reliable alternative method. The reverse predictive power, from WGS to FTIR, was slightly lower (AWC: 0.684, 95% CI: 0.406–0.962), reflecting the greater resolution of WGS. These findings suggest that while FTIR cannot fully replace WGS, it provides a reliable approximation of WGS-based classifications and a valuable first-line screening option or an alternative to WGS in settings where WGS is limited by cost or turnaround time.

### 4.6 Establishing FTIR-based classifier for *M. pachydermatis*

Using the IR Biotyper^®^ software, we trained an ANN model to classify haploid *M. pachydermatis* strains based on their FTIR spectra. The training set consisted of 139 spectra from 13 strains representing all three phylogroups, with multiple biological and technical replicates per strain to capture within-group variation (**Supplementary Figure S1)**. For validation, 153 spectra from 17 additional strains, also spanning all phylogroups, were withheld from training and used to assess model performance. After 142 training cycles, the model achieved 100% accuracy on the training set (**Supplementary Figure S8**).

Post validation, the automated classifier performance was evaluated using sensitivity, specificity, error rate, and predictive values. Validation results showed perfect classification for phylogroups I and II, with 108/108 spectra correctly identified, yielding 100% accuracy and sensitivity for these groups (**Table 3-4, Supplementary Table S6**). For phylogroup III, 43 out of 54 spectra were correctly classified (79.6% sensitivity), while the remaining 11 spectra from two strains were misclassified as either phylogroup I (n=5) or phylogroup II (n=6). These misclassifications lowered specificity for phylogroup I to 96.9% and phylogroup II to 96.3%, while specificity for phylogroup III remained at 100% (no false positives). Positive predictive values were high across groups—95.6% for phylogroup I, 94.7% for phylogroup II, and 100% for phylogroup III—as were negative predictive values (100% for phylogroups I and II, and 95.2% for phylogroup III).

**Table 3.** Confusion matrix showing classification performance of the IR Biotyper^®^ ANN model across *M. pachydermatis* phylogroups.

| Predicted / Actual | Phylogroup I | Phylogroup II | Phylogroup III |
| --- | --- | --- | --- |
| Phylogroup I | 108 | 0 | 5 |
| Phylogroup II | 0 | 108 | 6 |
| Phylogroup III | 0 | 0 | 43 |
| Misclassified * | 0 | 0 | 11 |

\* Misclassified spectra in phylogroup III, suggesting some spectral overlap with phylogroup I and II amongst some strains.

**Table 4.**
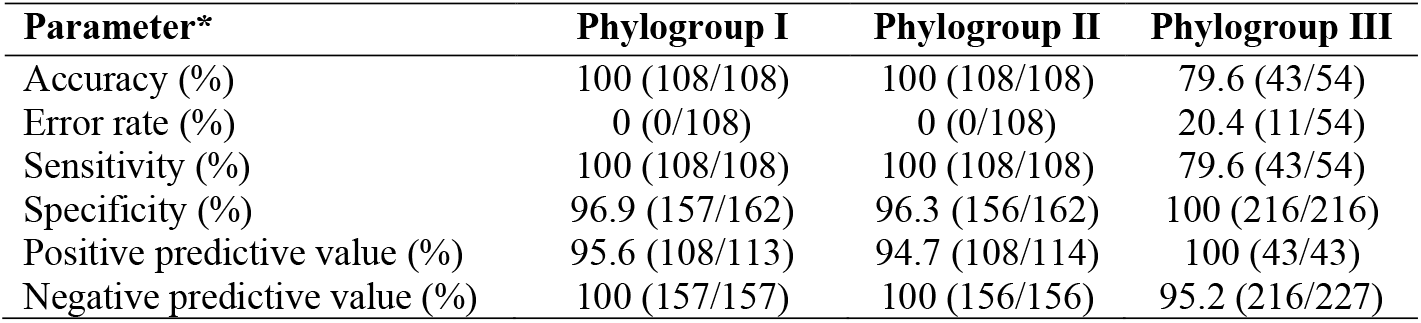
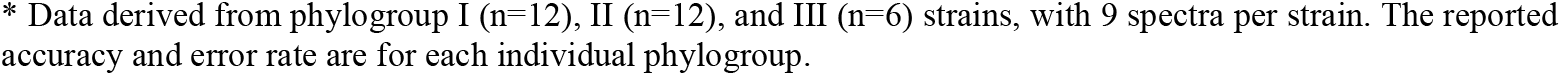
Classifier performance metrics per Phylogroup.

| Parameter* | Phylogroup I | Phylogroup II | Phylogroup III |
| --- | --- | --- | --- |
| Accuracy (%) | 100 (108/108) | 100 (108/108) | 79.6 (43/54) |
| Error rate (%) | 0 (0/108) | 0 (0/108) | 20.4 (11/54) |
| Sensitivity (%) | 100 (108/108) | 100 (108/108) | 79.6 (43/54) |
| Specificity (%) | 96.9 (157/162) | 96.3 (156/162) | 100 (216/216) |
| Positive predictive value (%) | 95.6 (108/113) | 94.7 (108/114) | 100 (43/43) |
| Negative predictive value (%) | 100 (157/157) | 100 (156/156) | 95.2 (216/227) |

\* Data derived from phylogroup I (n=12), II (n=12), and III (n=6) strains, with 9 spectra per strain. The reported accuracy and error rate are for each individual phylogroup.

Classifier confidence was further evaluated using the IR Biotyper^®^ traffic light color-coding system, which categorizes outlier values as green (<1.0, high confidence), yellow (1.0–2.0, moderate confidence), and red (>2.0, unclassifiable spectra). Of the 270 tested spectra, 188 (69.6%) received green scores and 82 (30.4%) received yellow scores (moderate confidence), with no spectra falling into the red category. Performance was similar for phylogroups I and II, while phylogroup III produced a higher proportion of yellow scores (**Figure 6**). Except for two misclassified strains from phylogroup III, the classifier consistently assigned green or yellow scores to all biological replicates of each strain. These misclassifications highlight the need for further refinement of the classifier for this subgroup; however, additional phylogroup III isolates were not available to us at the time of this study, preventing further optimization. Importantly, the classifier excluded all non-*M. pachydermatis* spectra (n=90) using the selected red score threshold, demonstrating its potential not only for intra-species strain typing but also for robust species-level exclusion. Overall, the model achieved sensitivity and accuracy of 95.9% (259/270), error rate 4.1% (11/270), and specificity of 98.0% (529/540) across all phylogroups. Together, these findings indicate that the FTIR-based classifier has strong potential for *M. pachydermatis* strain typing, broader diagnostic applications, and robust exclusion of unrelated species.

**Figure 6.**
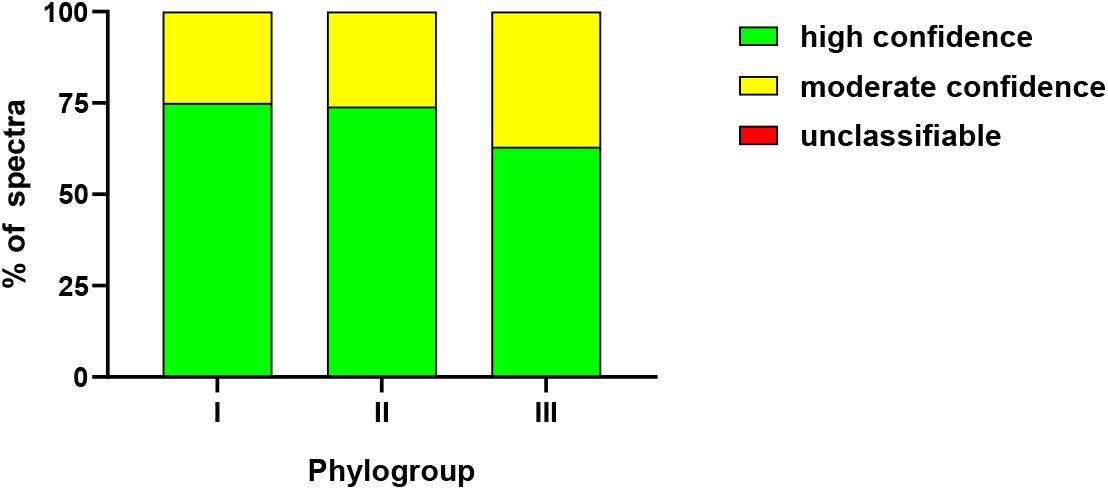
Classification confidence per phylogroup. The bar graph shows the percentage of spectra for each phylogroup, classified as green, yellow, or red, relative to the total number of spectra per phylogroup, from a minimum of nine individual absorption spectra (three independent biological replicates with three technical replicates each per strain). Classification scores are color-coded, green (<1.0) indicates spectra within the classifiers training set boundaries and considered reliable, yellow (1.0–2.0) represents spectra near the boundaries and considered moderately reliable, and red (>2.0) indicates spectra outside the training set boundaries, deemed unreliable.

## 5 Discussion

This study presents a novel FTIR spectroscopy protocol for analysing diversity in the skin commensal yeast *Malassezia* using the IR Biotyper^®^. Our findings expand upon and confirm an early report that used FTIR spectroscopy to differentiate *M. furfur, M. sympodialis, M. obtusa, M. slooffiae*, and *M. globosa* (42). Here, we validate FTIR spectroscopy as a reliable, rapid, and cost-effective alternative to WGS and to traditional molecular typing methods such as MLST for differentiating *Malassezia* species and strains in both epidemiological research and routine clinical or veterinary applications. The insights gained from this work enhance our understanding of *M. pachydermatis* diversity and population structure. In particular, WGS confirmed the existence of three well-supported phylogroups within *M. pachydermatis*, providing a robust genomic framework against which FTIR-based clustering could be evaluated.

The protocol we established for FTIR spectroscopy enables reproducible and consistent spectra acquisition. One of the challenges we faced during sample preparation was the poor ability to resuspend *Malassezia* cells in ethanol. We overcame this by first resuspending the fungal colony material in water and then adding ethanol, in contrast to the manufacturer’s standard “ethanol-first” protocol. Our observations suggest that *Malassezia* colonies exhibit high hydrophilicity, which may account for their improved resuspension in water. Overall, our protocol supported robust clustering of spectra acquired from biological replicates, in agreement with strain clustering by molecular methods.

The highest discriminatory power, at both the species and strain levels, was obtained from the spectral region corresponding to polysaccharides, suggesting that yeast cellular carbohydrates may vary significantly between *Malassezia* species and strains. This aligns with reported compositional differences in the carbohydrate-rich cell wall of *Malassezia* species (43). The carbohydrate-rich fungal cell wall, constituting of mannans, β-glucans, and chitin, is the primary interaction interface with the host and as such drives the host cell response to the fungus. The fungal cell wall also plays essential roles in fungal virulence and stress tolerance, with variations in its components influencing immune evasion, tissue adhesion, and biofilm formation. Notably, *Malassezia* cell walls contain abundant branched β-(1,6)-glucan and chitosan, which distinguishes it from the cell wall of other yeasts like *C. albicans* and *Saccharomyces cerevisiae* (44, 45). However, relatively little is known about inter- and intraspecies differences in *Malassezia* cell wall composition. Correlating specific FTIR spectral profiles with functional traits could therefore provide important insights into pathogenicity and immune stimulatory capacity and variations thereof across species and strains.

Growth conditions can also influence FTIR profiles by altering the micro- and macromolecular composition of cells. For instance, FTIR-based differentiation of *Pseudomonas aeruginosa* was superior when cells were grown on Mueller–Hinton agar compared to 5% sheep blood agar (46, 47), and greater reproducibility and discriminatory power was demonstrated for *Lactiplantibacillus plantarum* when cultured in liquid media rather than on agar (48). In *Malassezia*, media other than mDixon, the standard medium routinely used for culture and employed in our study, may alter the discriminatory power of FTIR spectroscopy. When considering alternative growth conditions for *Malassezia*, media simulating different skin environments, including healthy versus barrier-impaired and inflamed skin differing in pH, salt concentration, and lipid composition, should be explored (49–52). Because skin conditions are known to influence *Malassezia* growth and gene expression profiles (53, 54), they may also shape FTIR-detectable traits. This raises the possibility of applying FTIR to discriminate isolates from healthy versus diseased skin, which if causally linked to pathogenicity, could serve as a predictive tool for disease progression or clinical outcome in atopic dermatitis and other *Malassezia*-associated diseases. Our study further suggests that the FTIR approach has great potential if it is extended to other clinically important fungi in the future, such as *Cryptococcus*.

The *M. pachydermatis* strains included in our study are limited to haploid strains. In other *Malassezia* species, hybrid strains with chimeric genomes presumably arising from mating between haploid lineages, have been reported (32, 55). Hybridization can alter gene dosage, allele combinations, and regulatory networks, and may be accompanied by aneuploidy or segmental gene copy-number variation. Such changes could modify cell surface composition, lipid metabolism, and stress responses, all of which contribute to FTIR spectra. Whether hybrids occur in *M. pachydermatis*, and whether their FTIR signatures are distinguishable from those of haploid strains, remains to be determined. Addressing this will require confirmed hybrid isolates, matched culture conditions and growth phases, and joint analyses that relate FTIR clustering to whole-genome features such as ploidy, copy-number state, and parental haplotype composition. If hybrids are present and display distinct FTIR profiles, this would refine the classifier and further clarify how genomic context shapes the phenotypes captured by FTIR.

The development of an FTIR-based classifier for *M. pachydermatis* represents a significant advancement in the study of *Malassezia*, particularly given the limitations of traditional phenotypic identification methods. This approach provided valuable insights into within-species molecular diversity and phylogenetic relationships among strains, despite the modest training set. Expanding the training data to include a broader range of phylogenetically diverse strains will likely enhance the performance and applicability of the classifier across diverse settings. Looking ahead, FTIR may offer a rapid and accurate method for detecting potentially pathogenic *M. pachydermatis* strains, with implications for human and animal health. Realizing this potential will require further optimization and validation in multi-centre cohorts, as well as integration with WGS to identify the underlying genomic determinants of FTIR-detected phenotypic differences.

## 6 Acknowledgment

The authors would like to thank Philipp Bosshard for *M. restricta* (RS HS_19) and *M. globsa* (GB_AD72_270223) strains. We also thank Roger Stephan and members of the LeibundGut and Heitman labs for helpful advice and discussion. This work was supported by the Vetsuisse Faculty of University of Zurich and the National Institute of Health (NIH/NIAID R21 AI168672-02 to SLL and JH).

## 7 Author Contribution

**SK**: conceptualization, methodology, formal analysis, investigation, writing – original draft

**MDC**: formal analysis, data curation, writing – review & editing

**MDP**: resources, writing – review & editing

**LD**: resources, writing – review & editing

**GC**: resources, writing – review & editing

**JC**: resources, writing – review & editing

**JH**: resources, writing – review & editing

**SLL**: conceptualization, supervision, writing – review & editing, visualization, project administration, funding acquisition

**FM**: conceptualization, methodology, validation, formal analysis, investigation, writing -original draft, visualization, funding acquisition

